# A live attenuated influenza virus-vectored intranasal COVID-19 vaccine provides rapid, prolonged, and broad protection against SARS-CoV-2 infection

**DOI:** 10.1101/2021.11.13.468472

**Authors:** Junyu Chen, Pui Wang, Lunzhi Yuan, Liang Zhang, Limin Zhang, Hui Zhao, Congjie Chen, Yaode Chen, Jinle Han, Jizong Jia, Zhen Lu, Junping Hong, Liqiang Chen, Changfa Fan, Zicen Lu, Qian Wang, Rirong Chen, Minping Cai, Ruoyao Qi, Xijing Wang, Jian Ma, Min Zhou, Huan Yu, Chunlan Zhuang, Xiaohui Liu, Qiangyuan Han, Guosong Wang, Yingying Su, Quan Yuan, Tong Cheng, Ting Wu, Xiangzhong Ye, Changgui Li, Tianying Zhang, Jun Zhang, Huachen Zhu, Yixin Chen, Honglin Chen, Ningshao Xia

## Abstract

Remarkable progress has been made in developing intramuscular vaccines against severe acute respiratory syndrome coronavirus 2 (SARS-CoV-2); however, they are limited with respect to eliciting local immunity in the respiratory tract, which is the primary infection site for SARS-CoV-2. To overcome the limitations of intramuscular vaccines, we constructed a nasal vaccine candidate based on an influenza vector by inserting a gene encoding the receptor-binding domain (RBD) of the spike protein of SARS-CoV-2, named CA4-dNS1-nCoV-RBD (dNS1-RBD). A preclinical study showed that in hamsters challenged 1 day and 7 days after single-dose vaccination or 6 months after booster vaccination, dNS1-RBD largely mitigated lung pathology, with no loss of body weight, caused by either the prototype-like strain or beta variant of SARS-CoV-2. Lasted data showed that the animals could be well protected against beta variant challenge 9 months after vaccination. Notably, the weight loss and lung pathological changes of hamsters could still be significantly reduced when the hamster was vaccinated 24 h after challenge. Moreover, such cellular immunity is relatively unimpaired for the most concerning SARS-CoV-2 variants. The protective immune mechanism of dNS1-RBD could be attributed to the innate immune response in the nasal epithelium, local RBD-specific T cell response in the lung, and RBD-specific IgA and IgG response. Thus, this study demonstrates that the intranasally delivered dNS1-RBD vaccine candidate may offer an important addition to fight against the ongoing COVID-19 pandemic, compensating limitations of current intramuscular vaccines, particularly at the start of an outbreak.

## Introduction

Coronavirus disease 2019 (COVID-19), caused by severe acute respiratory syndrome coronavirus 2 (SARS-CoV-2), has had an immeasurable impact on health, the economy and social stability worldwide^1,2^. The rapid development of multiple COVID-19 vaccines has been an incredible scientific achievement^3^. Multiple vaccines based on traditional or modern platform technologies have demonstrated high effectiveness for preventing severe COVID-19, hospitalization and death in clinical trials as well as in the real world for at least several months^4–8^, enabling widespread vaccine administration to curb the COVID-19 pandemic globally. Nonetheless, the effectiveness of current vaccines in interrupting human-to-human transmission and for mild or asymptomatic patients has been well below expectations, especially for variants with stronger transmissibility and antigenic changes, such as the delta variant. Indeed, the number of newly confirmed cases is increasing rapidly again even in countries with extremely high levels of vaccine coverage^9–13^. Thus, it is imperative to continue developing new COVID-19 vaccines using different vaccine strategies.

To date, COVID-19 vaccines approved for use by different regulatory authorities, including mRNA vaccines, inactivated vaccines, recombinant adenovirus vaccines and recombinant protein vaccines, are all administered through traditional muscle injection, which are commonly limited for their ability to induce mucosal immunity and local immunity^14–17^. While some countries with sufficient vaccine supplies have been achieving the potential “herd” immunity ^18^, breakthrough infections are common among vaccinated people. Importantly, majority of children are not among the vaccinated groups. With countries reopening borders for international travelers and the increasing emergence of variants of concern, epidemics with high transmission among specific groups of people will become very common. Solutions in response to the evolving COVID-19 pandemic are imminently needed. Given the predominant respiratory tropism of SARS-CoV-2 and the evidence that intranasal live attenuated influenza vaccine (LAIV) has equivalent and even improved efficacy compared with that of inactivated influenza vaccine (IIV) ^19,20^, several vaccine candidates intended to be delivered by intranasal administration or inhalation are under development, and some of them have shown potential in animal models and early phase clinical trials ^21,22^. To our knowledge, eight intranasally delivered COVID-19 vaccines have been tested in clinical trials globally, seven of which are based on virus vectors, including adenovirus, respiratory syncytial virus and influenza virus^15,23,24^. These intranasal vaccines have shown the potential to elicit mucosal IgA and CD8^+^ T cell-mediated immune responses in the respiratory tract as well as serum IgG responses, resulting in more efficiently reduction of virus replication and shedding in both the lungs and the nasal passages than intramuscular vaccination ^15,25,26^.

Here, we present data demonstrating the rapid (1 day), prolonged (9 months) and broad protection of and comprehensive innate and adaptive immune responses to an intranasally delivered COVID-19 vaccine based on a live attenuated influenza virus vector in animal models. This vaccine candidate has been shown to be well tolerated and immunogenic in Chinese adults, and a global multicenter phase III clinical trial will be initiated soon.

## Results

### Construction and pathogenic analysis of the dNS1-RBD vaccine candidate

The vaccine candidate CA4-dNS1-nCoV-RBD (dNS1-RBD) was constructed by inserting a gene encoding the receptor-binding domain (RBD) of the spike protein of the SARS-CoV-2 prototype strain into the previously reported NS1-deleted backbone of H1N1 influenza virus California/4/2009 (CA04-dNS1)^27^ (Fig. 1a). We compared the growth kinetics of dNS1-RBD with those of the wild-type A/California/04/2009(H1N1) parental virus (CA04-WT) and its NS1-deleted version (CA04-dNS1) in Madin-Darby canine kidney (MDCK) cells. As expected, the replication of dNS1-RBD was significantly suppressed at 37°C and 39°C compared with that at 33°C due to the existence of temperature-sensitive mutations in the CA04-dNS1 vector (Fig. 1b), which is a desirable feature for reducing the risk of influenza-associated adverse reactions in the lung. In line with the above results, at seven days post nasal inoculation, all six ferrets in the CA04-WT group showed viral shedding in the nasal turbinate and throat, in contrast to none of the ten ferrets in the dNS1-RBD group (Supplementary Fig. S1). The expression of the RBD and HA antigen in dNS1-RBD-infected MDCK cells was visualized using confocal analysis and further confirmed by Western blot (Fig. 1c, d). Evaluated by ten continuous passages, the genetic and expression stability of the RBD fragment of dNS1-RBD in the MDCK cell culture system seemed acceptable for large-scale production (Fig. 1e).

**Fig. 1.**
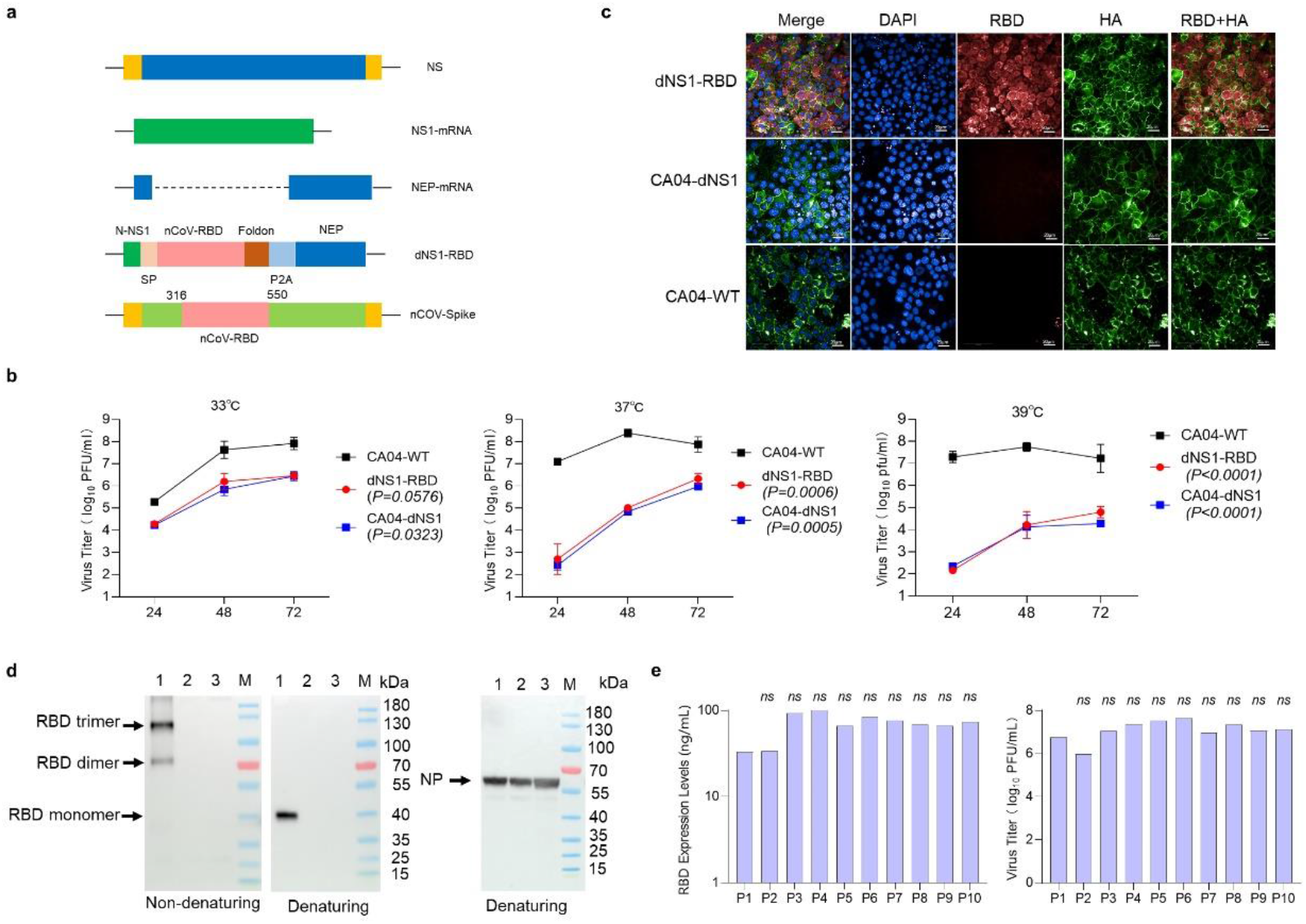
Construction and characterization of a recombinant live attenuated influenza virus-based SARS-CoV-2 vaccine. **a** Construction of an mRNA-encoding plasmid that transcribes DelNS1 with RBD-inserted mRNA. RBD, receptor-binding domain. **b** Replication efficiency of the dNS1-RBD, CA04-dNS1 and CA04-WT viruses varied with 33°C , 37°C and 39°C conditions in MDCK cells. Data represent the mean values ± SDs of results from three independent experiments. Analysis was performed by two-way repeated-measures analysis of variance (ANOVA). Significance was set at p <0.05. **c** Confocal analysis of the RBD and HA protein expressed by the influenza vector in MDCK cells. The coexpression of RBD and HA could be detected only for dNS1-RBD. MDCK cells were fixed 72 h after infection. Red fluorescence indicates the RBD; green fluorescence indicates HA. **d** Immunoblot analysis of RBD and NP expression in denatured and nondenatured cell lysate samples 36 h after infection by dNS1-RBD (1), CA04-dNS1 (2) and CA04-WT (3). Most of the secreted RBD protein for dNS1-RBD formed an RBD trimer, with RBD rarely existing in the dimer form. **e** Plaque assay and sandwich ELISA analysis of RBD expression was performed on the virus supernatant harvested from serial passages 1 to 10 of dNS1-RBD. ns, not significant (P > 0.05). Significance was determined by one-way ANOVA with the Kruskal-Wallis test.

Intranasal inoculation in BALB/c mice and ferrets confirmed the obvious attenuation of dNS1-RBD compared to the parental CA04-WT virus (Supplementary Fig. S2). Mice inoculated with 10^5^-10^7^ plaque-forming units (PFU) of the parental CA04-WT virus succumbed to infection after seven days, whereas mice inoculated with dNS1-RBD continued to maintain their weight. Likewise, for ferrets, which are highly susceptible to influenza virus infection, inoculation with 10^7^ PFU of CA04-WT but not dNS1-RBD resulted in obvious influenza-like symptoms, with fever, weight loss and pathological injury in lung tissues. In summary, a recombinant live attenuated influenza virus stably expressing the SARS-CoV-2 RBD segment with remarkably less virulence than its parental influenza virus was generated.

### Rapid and prolonged protection against infection with the prototype and beta variant of SARS-CoV-2 in hamsters after intranasal immunization with dNS1-RBD

The expected dominant advantage of intranasal immunization is the establishment of an immune barrier in the respiratory tract, which is particularly desired for prevention of respiratory virus infection. Hence, to test the protective effects of dNS1-RBD at 1 day or 7 days after single-dose immunization and 6 months after two doses of dNS1-RBD (prime and boost regimen with a 14-day interval), we chose the interanimal transmission model in golden Syrian hamsters to mimic the predominant natural route of SARS-CoV-2 infection (Fig. 2a). The model is preferred because it has been demonstrated to be sensitive to SARS-CoV-2 infection and associated COVID-19-like lung damage and can support efficient viral transmission from inoculated hamsters to naïve hamsters by direct contact and via aerosols ^28,29^. Vaccinated or sham hamsters were infected through cohousing with donor hamsters infected by the prototype strain or the beta variant. The sham hamsters showed continuous body weight loss beginning 2 days post infection (dpi), with maximal weight loss at 7 dpi (mean: -9.7% for the prototype virus challenge group and -12.2% for the beta variant challenge group); in contrast, weight loss was not obvious in animals of all vaccine groups (Fig. 2b, c). Lung damage at 5 dpi was quantitatively measured using a comprehensive pathological scoring system. Animals in the sham groups had significantly higher pathological scores than those in the vaccine groups (Fig. 2d, e). The pathological histology analysis of lung tissues (Fig. 2f) and gross lung images (Supplementary Fig. S3) taken at 5 dpi showed that vaccinated hamsters were largely protected from lung damage caused by infection with the SARS-CoV-2 prototype strain and the beta variant, with minimal, if any, focal histopathological changes in the lung lobes. In contrast, hamsters in unvaccinated groups developed severe lung pathology with consolidated pathological lesions and severe or intensive interstitial pneumonia characterized by inflammatory cell infiltration in a focally diffuse or multifocal distribution. On average, 30% to 50% of the alveolar septa of these unvaccinated animals became thicker, resembling findings in patients with severe COVID-19 bronchopneumonia ^28,29^. Additionally, at 5 dpi, the viral loads in the lung tissue of vaccinated hamsters, except hamsters in group 2 and group 3, which were challenged with the prototype virus at 6 months after receiving two doses of vaccine, were significantly lower than those in the lung tissue of sham controls (Supplementary Fig. S4). Best of all, the immunized hamsters were protected against the beta variant challenge at 9 months after two-dose vaccination (Fig. 3a), with a reduction of 1.5 log10 in viral RNA loads (Fig. 3b), and of 5.1 log10 for comprehensive pathological scores (Fig. 3c) as compared to sham group. Moreover, the lungs of vaccinated animals remained normal, or near to normal with no more signs of bronchopneumonia (Fig. 3d).

**Fig. 2.**
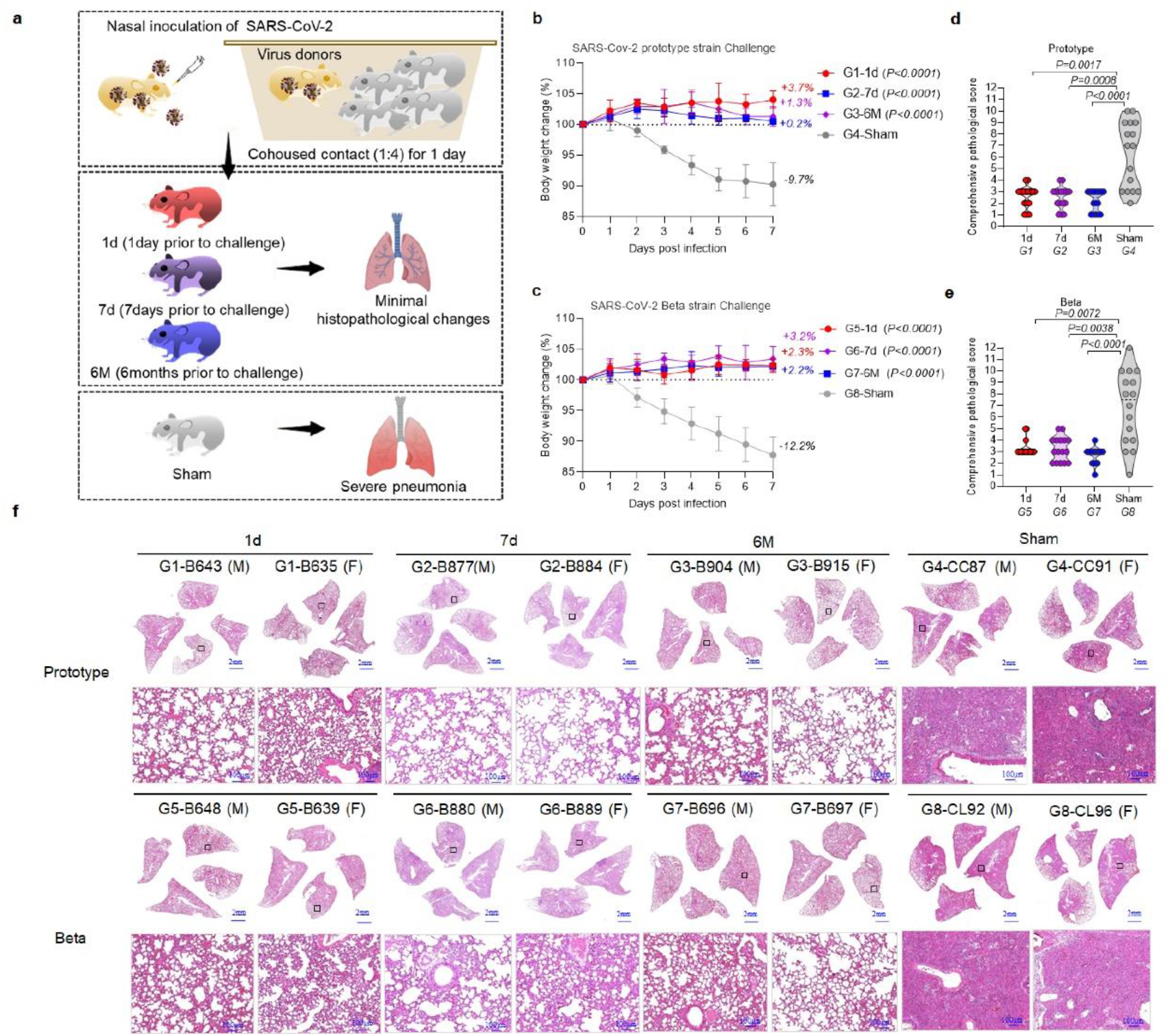
Protective efficacy of dNS1-RBD against SARS-CoV-2 challenge in Syrian hamsters one day, seven days and 6 months after vaccination. **a** Vaccination and challenge scheme. There were eight experimental groups, each containing eight hamsters (male:female =1:1). Groups 1, 2, 5 and 6 received a single dose of dNS1-RBD 1 day or 7 days before virus challenge; groups 3 and 7 received two doses of dNS1-RBD at a 14-day interval 6 months before virus challenge; and groups 4 and 8 served as sham controls and were not treated. Every 4 vaccinated or sham control hamsters were cohoused with 1 donor animal for 24 h and then observed for 7 days of follow-up. **b-c** Changes in the body weights of hamsters were recorded following cohousing exposure. The average weight loss of each group at 7 dpi is indicated as a colored number. Data shown are the mean ± SD. Significant differences compared to the sham group were analyzed using two-way repeated-measures analysis of variance (ANOVA) with Dunnett’s multiple comparisons test. **d-e** Comprehensive pathological scores of the hamster lungs. Scores were determined based on the severity and percentage of injured areas for each lung lobe collected from the indicated animal. Significance compared to the sham group was determined by one-way ANOVA with the Kruskal-Wallis test. **f** H&E staining of lung sections from tested hamsters collected on day 5 after cohousing exposure. Views of the whole lung lobes (4 independent sections) are presented in the above panel, and the areas in the small black boxes are enlarged in the lower panel.

**Fig. 3.**
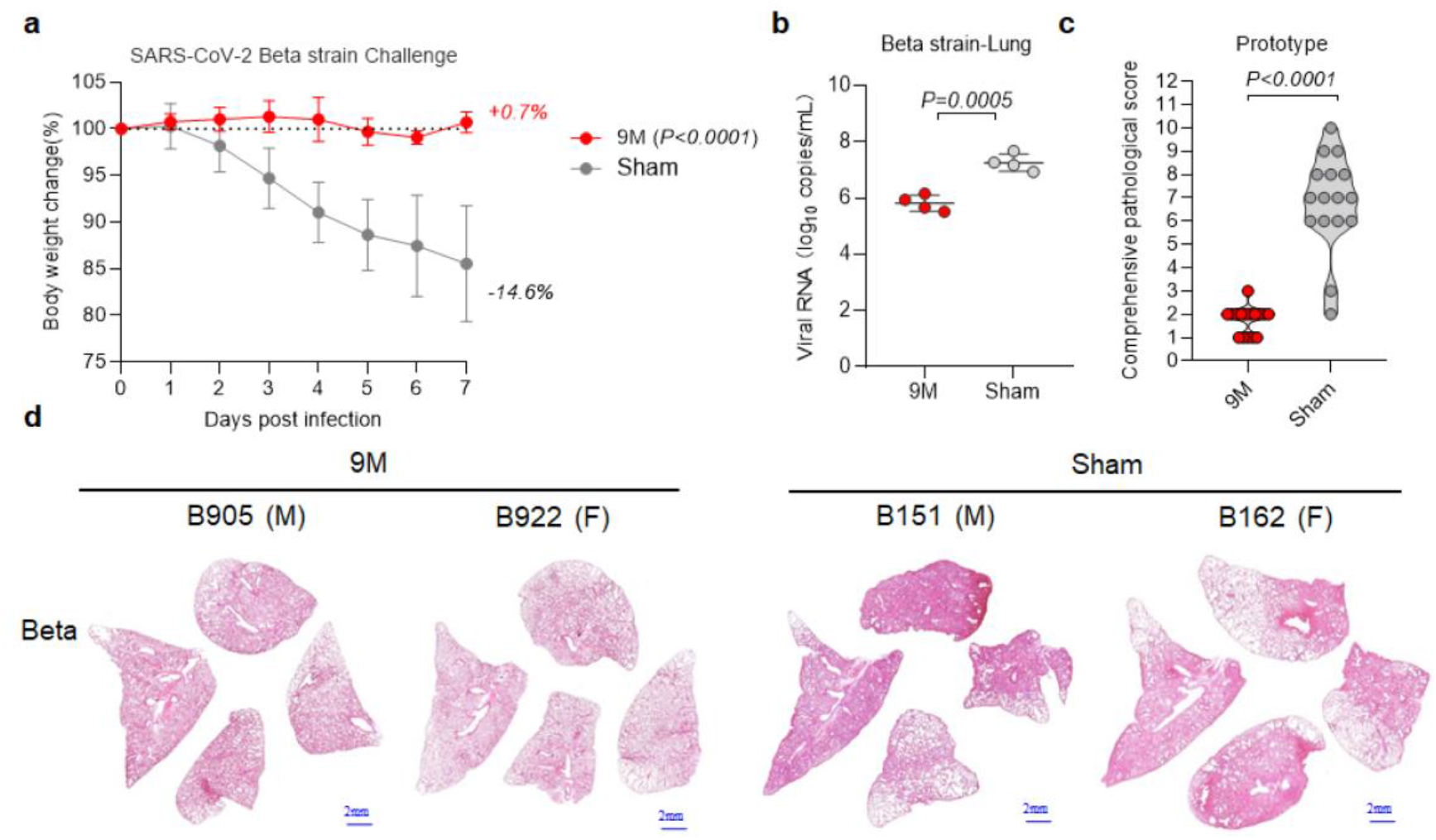
Protective efficacy of dNS1-RBD against SARS-CoV-2 challenge in Syrian hamsters 9 months after vaccination. **a** Changes in the body weights of hamsters were recorded following cohousing exposure. The average weight loss of each group at 7 dpi is indicated as a colored number. Data shown are the mean ± SD. Significant differences compared to the sham group were analyzed using two-way repeated-measures analysis of variance (ANOVA) with Dunnett’s multiple comparisons test. **b** Viral loads of lung tissue obtained at 5 dpi in hamsters challenged post inoculation were determined by RT-PCR. Data are the mean ± SD; significance was determined by two-tailed Student’s t-test. Symbols represent individual hamsters. **c** Comprehensive pathological score of hamster lung pathology images. Scores were determined based on the severity and percentage of injured areas for each lung lobe collected from the indicated animal. Significance was determined by a two-tailed Mann-Whitney U test. **d** H&E staining of lung sections from tested hamsters collected on day 5 after cohousing exposure. Views of the whole lung lobes (4 independent sections) are presented.

Taken together, these results demonstrated that dNS1-RBD vaccination could efficiently block the pathogenicity of homogeneous and heterogeneous SARS-CoV-2 infection in golden Syrian hamsters in the direct contact model in the short term and long term.

The above rapid and robust protection conferred by dNS1-RBD encouraged us to explore the protective effects of the vaccine candidate with postexposure immunization. After 24 h of cohousing with donors infected by the prototype strain, the infected hamsters were inoculated with a single dose of dNS1-RBD (Fig. 4a). The sham hamsters showed continuous body weight loss, with maximal weight loss at 7 dpi (mean: -10.0%); in contrast, the weight loss was reduced in animals in the vaccinated group (mean: -5.5%) (Fig. 4b). Although the viral loads in the lung tissue of vaccinated hamsters were not reduced compared to those in the lung tissue of the sham group hamsters (Fig. 4c), lung damage was significantly mitigated (Fig. 4d, e).

**Fig. 4.**
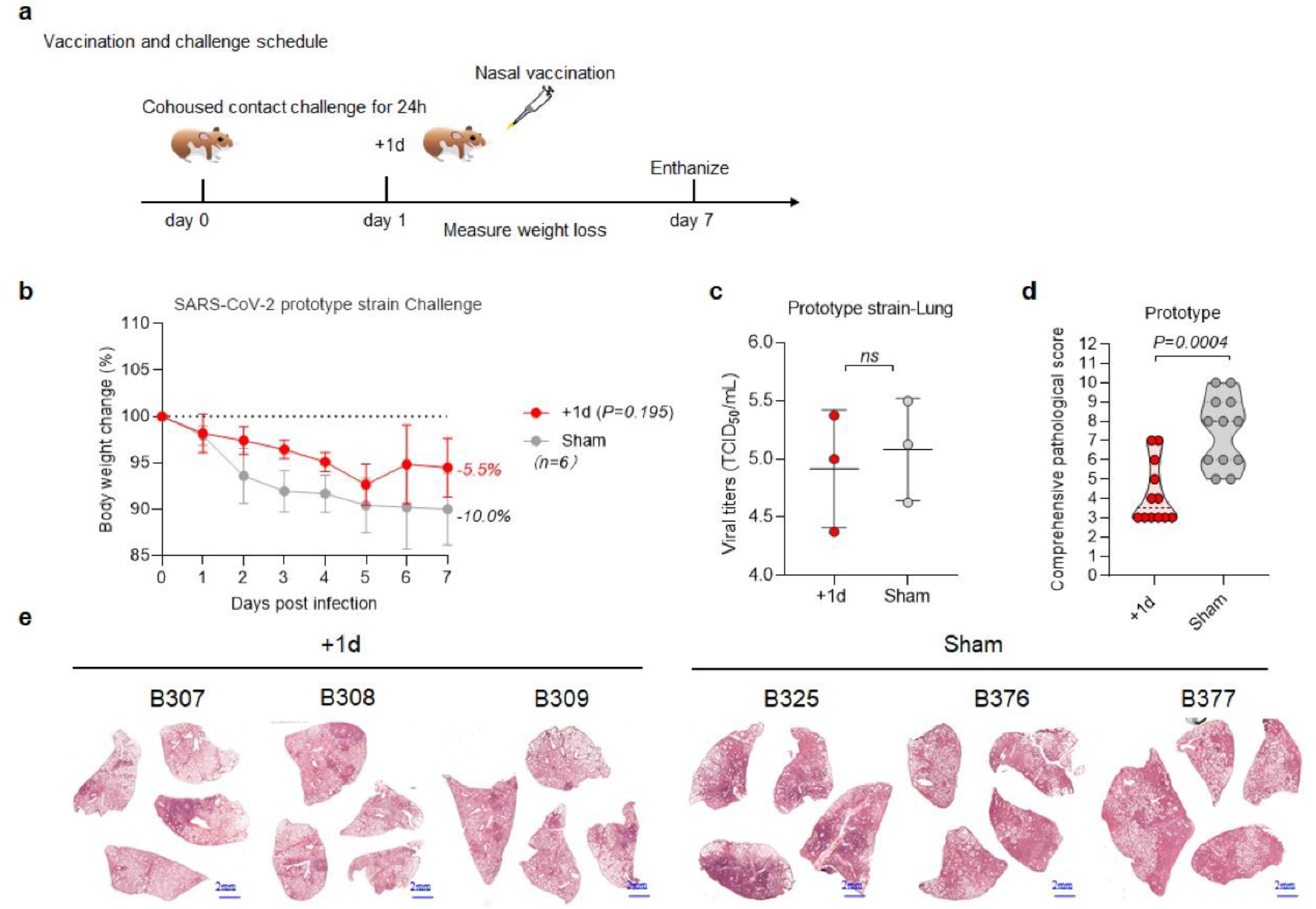
Protective efficacy of dNS1-RBD against SARS-CoV-2 challenge in Syrian hamsters 1 day before vaccination. **a** Vaccination and challenge scheme. Every 4 vaccinated or sham control hamsters were cohoused with 1 donor animal for 24 h. After 24 h of cohousing, the vaccinated hamsters were inoculated with a single dose of dNS1-RBD. Each animal was measured daily for body weight before sacrifice. **b** Changes in the body weights of hamsters were recorded following cohousing exposure. The average weight loss of each group at 7 dpi is indicated as a colored number. Data shown are the mean ± SD. Significant differences compared to the sham group were analyzed using two-way repeated-measures analysis of variance (ANOVA) with Dunnett’s multiple comparisons test. **c** Viral loads of lung tissue obtained at 5 dpi in hamsters challenged post inoculation were determined by TCID_50_ assay. Data are the mean ± SD; ns, not significant (P > 0.05); significance was determined by two-tailed Student’s t-test. Symbols represent individual hamsters. **d** Comprehensive pathological score of hamster lung pathology images. Scores were determined based on the severity and percentage of injured areas for each lung lobe collected from the indicated animal. Significance was determined by a two-tailed Mann-Whitney U test. **e** H&E staining of lung sections from tested hamsters collected on day 5 after cohousing exposure. Views of the whole lung lobes (4 independent sections) are presented.

### Protection against prototype SARS-CoV-2 infection in hACE2-humanized mice after intranasal immunization with dNS1-RBD

Although dNS1-RBD conferred rapid and lasting protection in the hamster model, only a weak IgG response could be detected in vaccinated hamsters (Supplementary Fig. S5), and the lack of a reliable detection system for measuring mucosal IgA, the T cell immune response and innate immune biomarkers hampered efforts to understand the vaccine-induced protective immune mechanism. Hence, we chose a mouse model to understand the mechanism of the rapid and lasting robust protective activity provided by intranasal immunization using dNS1-RBD. First, we validated the efficacy of intranasal immunization with dNS1-RBD in hACE2-humanized mice and then mapped the profile of innate and adaptive immune responses in BALB/c and C57BL/6 mice. Previous studies have demonstrated that hACE2-humanized mice created by CRISPR/Cas9 knock-in technology are susceptible to SARS-CoV-2 infection upon intranasal inoculation, and the resulting pulmonary infection and pathological changes resemble those observed in COVID-19 patients^30^. We evaluated the immunogenicity and protective effects of the dNS1-RBD vaccine candidate in hACE2-KI/NIFDC mice. All mice were immunized twice through the intranasal route at day 0 and day 14. On day 28 (14 days after the second immunization), the vaccinated group and control group were intranasally challenged with 1×10^4^ PFU SARS-CoV-2 per mouse under anesthesia (Fig. 5a). Compared to the severe weight loss of mice in the control group post infection, the weight change of mice in the vaccinated group was essentially negligible (Fig. 5b). At day 14, all vaccinated hACE2-KI/NIFDC mice showed a moderate level of RBD-specific IgG (Fig. 5c). We next determined the viral loads in the lung tissue by RT-PCR and plaque assay after all mice were euthanized at 4 dpi. All sham-treated mice had a high viral load (mean 10^5.12^ copies/mL and 10^4.66^ PFU/mL) at 4 dpi. In contrast, the viral load in the lung tissue of the vaccinated mice significantly decreased to a mean of 10^3.99^ copies/mL and 10^2.47^ PFU/mL (Fig. 5d). As expected, all mice in the sham group developed severe interstitial pneumonia characterized by inflammatory cell infiltration and alveolar septal thickening. In contrast, all vaccinated mice were largely protected from the damage caused by SARS-CoV-2 infection, with very mild and focal histopathological changes in a few lobes of the lung (Fig. 5e). Overall, dNS1-RBD vaccination efficiently limited the pathogenicity of SARS-CoV-2 infection in hACE2-humanized mice.

**Fig. 5.**
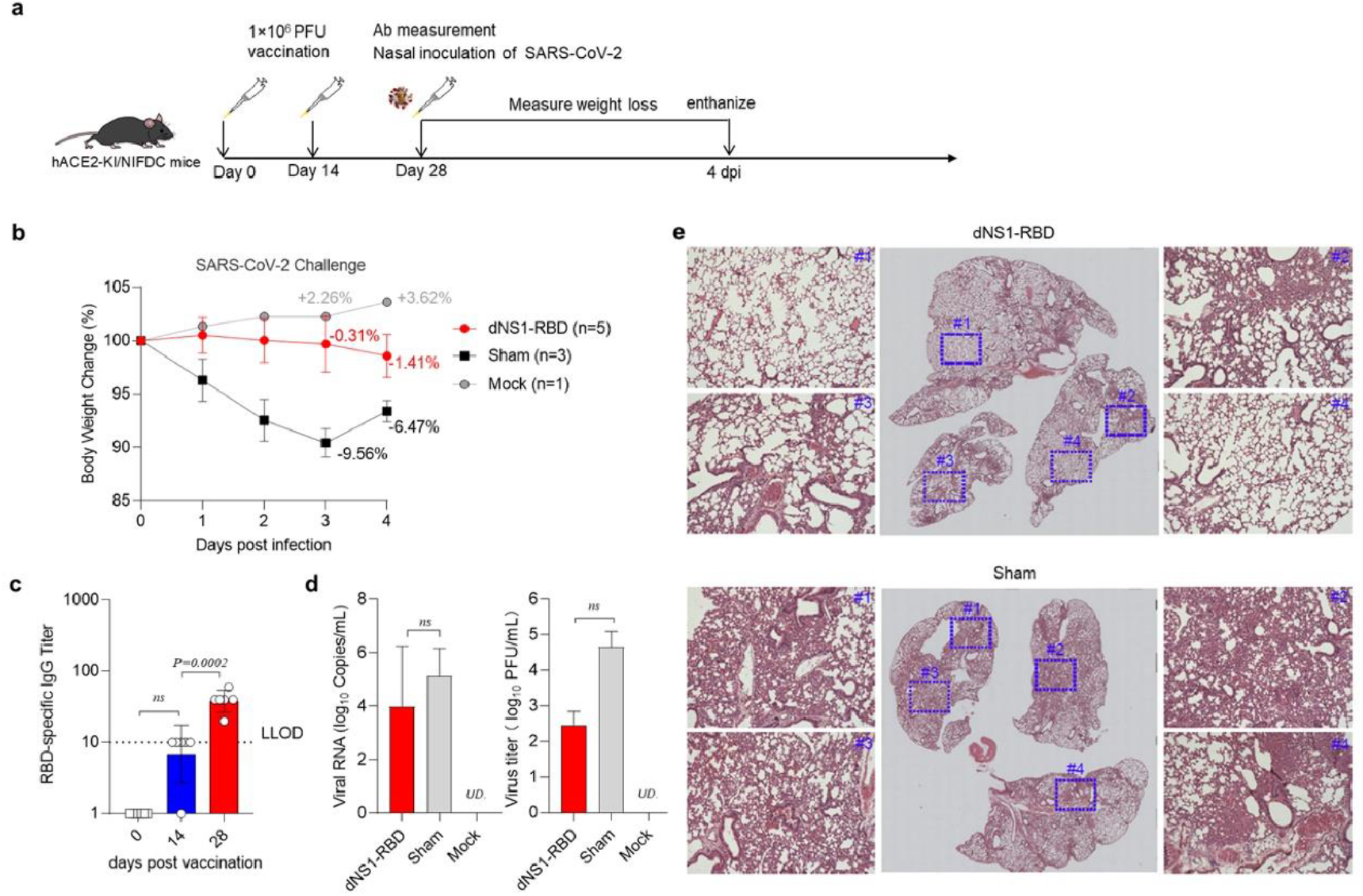
Immunogenicity and protective efficacy of dNS1-RBD against SARS-CoV-2 in hACE2-KI/NIFDC mice. **a** Experimental strategy. hACE2-KI/NIFDC mice were immunized twice at an interval of 14 days through the intranasal route with 1×10^6^ PFU of dNS1-RBD (n=5). **b-e** The protective efficacy of dNS1-RBD against SARS-CoV-2 challenge at 14 days after the second immunization was evaluated in hACE2-KI/NIFDC mice. **b** Changes in the body weights of mice were recorded. **c** The RBD-specific IgG response on the indicated days after the first vaccination was detected. LLOD-lower limit of detection. Data for antibody analysis are presented as the geometric mean with the geometric SD from four independent experiments; ns, not significant (P > 0.05). Significance compared to samples collected at day 0 was determined by ordinary one-way ANOVA multiple comparison. Symbols represent individual animals. **d** Viral loads in lung tissue obtained at 4 days post inoculation were determined by RT-PCR and plaque assay. Data are presented as the mean ± SD from four independent experiments; error bars reflect the SD. ns, not significant (P > 0.05); two-tailed Student’s t-test was used for intergroup statistical comparison. **e** Histopathological examinations of lungs from dNS1-RBD-immunized A53 mice and unvaccinated sham control A69 mice at day 4 post infection. The areas in the small box colored in blue were enlarged around the whole lung lobes.

### Intranasal inoculation of dNS1-RBD promotes comprehensive local immunity in the respiratory tract

It is well recognized that at least several days or weeks are needed before protective adaptive immunity is adequately activated. To understand the mechanism of the rapid and robust protection induced by intranasal administration of dNS1-RBD, the levels of innate immune response biomarkers in the respiratory tract of BALB/c mice after intranasal administration of dNS1-RBD were compared to those in unvaccinated controls and animals infected with wild-type influenza virus CA04-WT (Fig. 6a and Supplementary Fig. S6). The levels of the proinflammatory cytokines and chemokines IL-6, IL-18, IFN-γ, IFN-α, MCP-1, IP-10, MIP-1α, and MIP-1β, which are linked to the activation of innate immunity against respiratory viruses, were significantly elevated in lung tissue of mice 24 h post immunization. Simultaneously, the activation of myeloid dendritic cells (DCs) in the spleen and macrophages in cervical lymph nodes was observed in vaccinated mice 14 days post immunization (Supplementary Fig. S7), which have been reported to be associated with innate immunity with memory characteristics, i.e., trained immunity ^31^. As they were treated with a nasal spray vaccine, dNS1-RBD-vaccinated animals were expected to produce robust cellular-mediated immunity (CMI) after prime-boost immunization and have a significantly greater number of RBD-specific immune cells within the respiratory system than among peripheral blood mononuclear cells (PBMCs) or lymphocytes from the spleen and cervical lymph nodes (Fig. 6b), which suggested that the CMI response induced by dNS1-RBD is local and intensive in the respiratory tract. In particular, the RBD-specific cellular immune response was 22 times higher than that in PBMCs (Fig. 6b), which poses a challenge in evaluating the immune response of this vaccine based on PBMC test results in clinical trials.

**Fig. 6.**
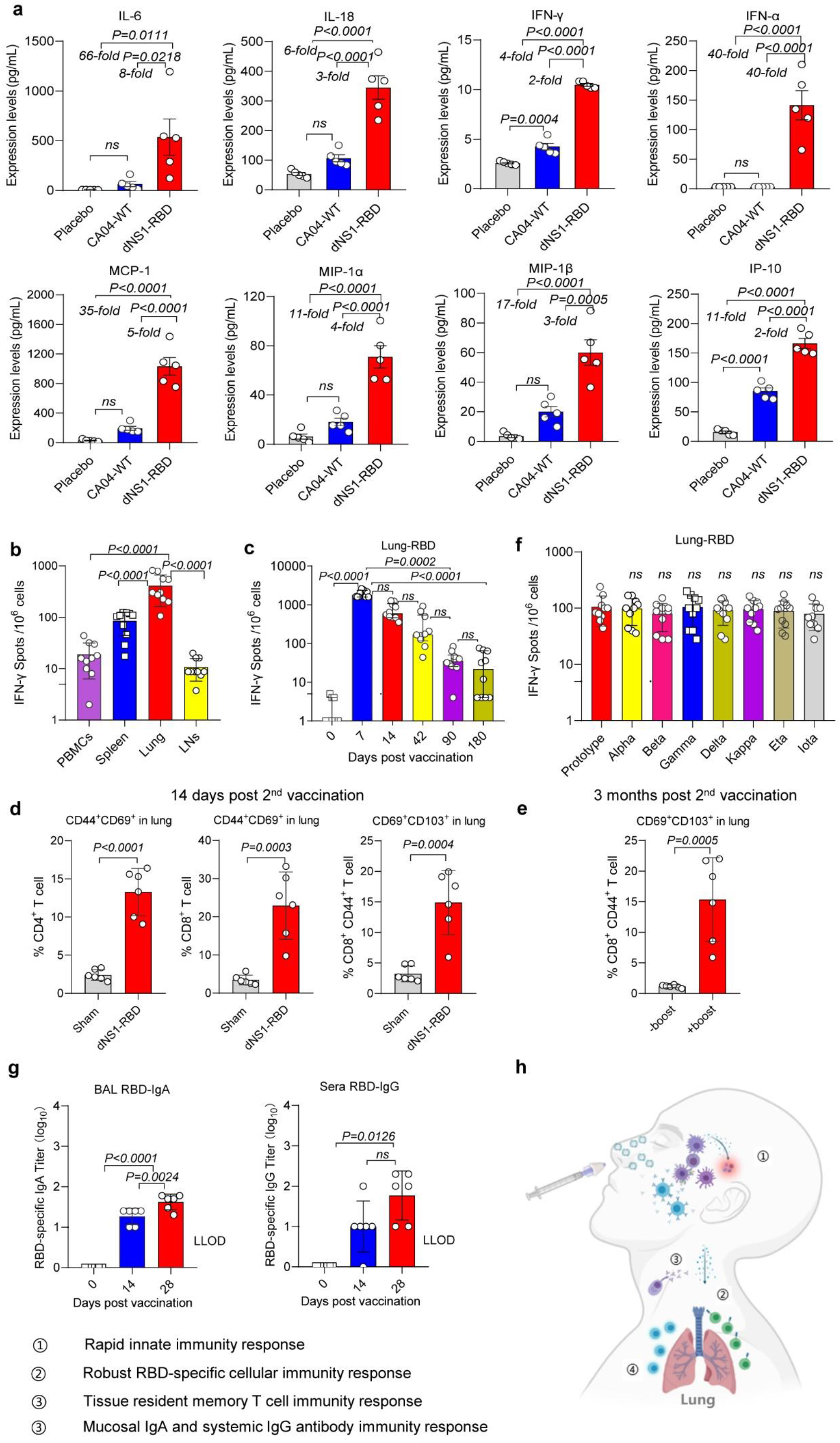
The profile of immune responses in the respiratory tract and blood induced by intranasal administration of dNS1-RBD in mice. **a** Lung homogenates from BALB/c mice vaccinated with dNS1-RBD one day prior to sacrifice were assayed for cytokine and chemokine expression levels by ProcartaPlex immunoassays. The data are expressed as ng/mL. **b** Lymphocytes in the peripheral blood, spleen, lung and lymph nodes collected 14 days after booster immunization at day 14 were subjected to IFN-γ ELISPOT assays. **c** IFN-γ ELISPOT assays for pulmonary lymphocytes from C57BL/6 mice vaccinated intranasally with two doses, with a booster at day 14 to assess T cell response kinetics at the indicated time points. Data are the median with IQR; ns, not significant (P > 0.05); significance was determined by one-way ANOVA with the Kruskal-Wallis test. **d** TRM markers expressed in pulmonary T cells 2 weeks after booster vaccination at a 14-day interval. **e** TRM markers expressed in pulmonary T cells were detected in mice boosted with an extra 3rd inoculation 3 months after booster vaccination with a 14-day interval. **f** The responses were assessed under prestimulation of various peptide pools covering the spike of SARS-CoV-2 variants. **g** RBD-specific IgA levels in BAL fluid and IgG levels in serum were measured by ELISA for BALB/c mice vaccinated twice at day 0 and day 14. Data for antibody analysis are presented as the geometric mean with the geometric SD from four independent experiments. LLOD-lower limit of detection. **h** Schematic overview of the immune response to the dNS1-RBD vaccine in the respiratory tract. Data are the mean ± SD; ns, not significant (P > 0.05); significance was determined by ordinary one-way ANOVA multiple comparison (**a**, **b**, **d** and **f**) or two-tailed Student’s t-test (**e**).

For RBD-specific T cell activation and proliferation, the CMI response reached a peak at 7 days after a single-dose intranasal administration, with more rapid and robust response dynamics compared to those of the humoral response, and fell to a moderate level at 42 days following the prime-boost regimen with a 2-week interval (Fig. 6c). Although the CMI response progressively waned, the specific T cell response from 9/10 animals was detectable at 3 months by IFN-γ ELISpot after booster immunization, with 6/10 animals further proven to be positive at 6 months. In addition to the longevity of vaccine-induced immunity, a substantial number of CD4^+^ and CD8^+^ T cells in the lungs of mice vaccinated with dNS1-RBD showed upregulated expression of the TRM marker CD69, while dNS1-RBD-generated CD8^+^ TRM cells also expressed the canonical CD8^+^ TRM marker CD103 (Figs. 6d and Supplementary Fig. S8), indicating that vaccination with dNS1-RBD generated lung-resident memory RBD-specific CD4^+^ and CD8^+^ TRM populations. Three months post 2^nd^ vaccination with a 14-day interval, activation and proliferation of memory CD69^+^CD103^+^ TRM cells could be detected 7 days after the boost inoculation (Fig. 6e).

As a recent study showed that SARS-CoV-2 variants (B.1.1.7 in the UK, B.1.351 in South Africa, B.1.525 in Nigeria and P1 in Brazil) are relatively resistant to serum from individuals who have recovered from COVID-19 or serum from individuals who have been vaccinated against SARS-CoV-2^32^, we used peptides covering the RBD with key mutations from the major variants (including alpha, beta, gamma, delta, kappa, eta, and iota) and prototype strains to stimulate lymphocytes and found similar RBD-specific T cell responses in the lungs from vaccinated mice, suggesting that the key mutants are still covered by the dNS1-RBD vaccine (Fig. 6f).

In-depth profiling of the T cell compartment by intracellular cytokine staining confirmed a significant increase in RBD-specific IFN-γ^+^ effector memory T cells in the lung, spleen and cervical lymph nodes (Fig. 7b), RBD-specific TNF-α^+^ CD8^+^ T cells in the lung and spleen (Fig. 7c) and RBD-specific IL-2^+^CD8^+^ T cells in the lung (Fig. 7d) from immunized mice in comparison with those from mice in the control group upon *ex vivo* stimulation with pools of overlapping 15-mer RBD peptides. A significant enrichment of other subpopulations, such as IL-2^+^, IFN-γ^+^ and TNF-α-expressing CD4^+^ T lymphocytes, was not observed (data not shown). The robust production of IFN-γ from CD8^+^ T cells indicated a favorable immune response with both antiviral and immune-augmenting properties, suggesting the induction of a Th1-biased cellular immune response and the potential safety of this vaccine.

**Fig. 7.**
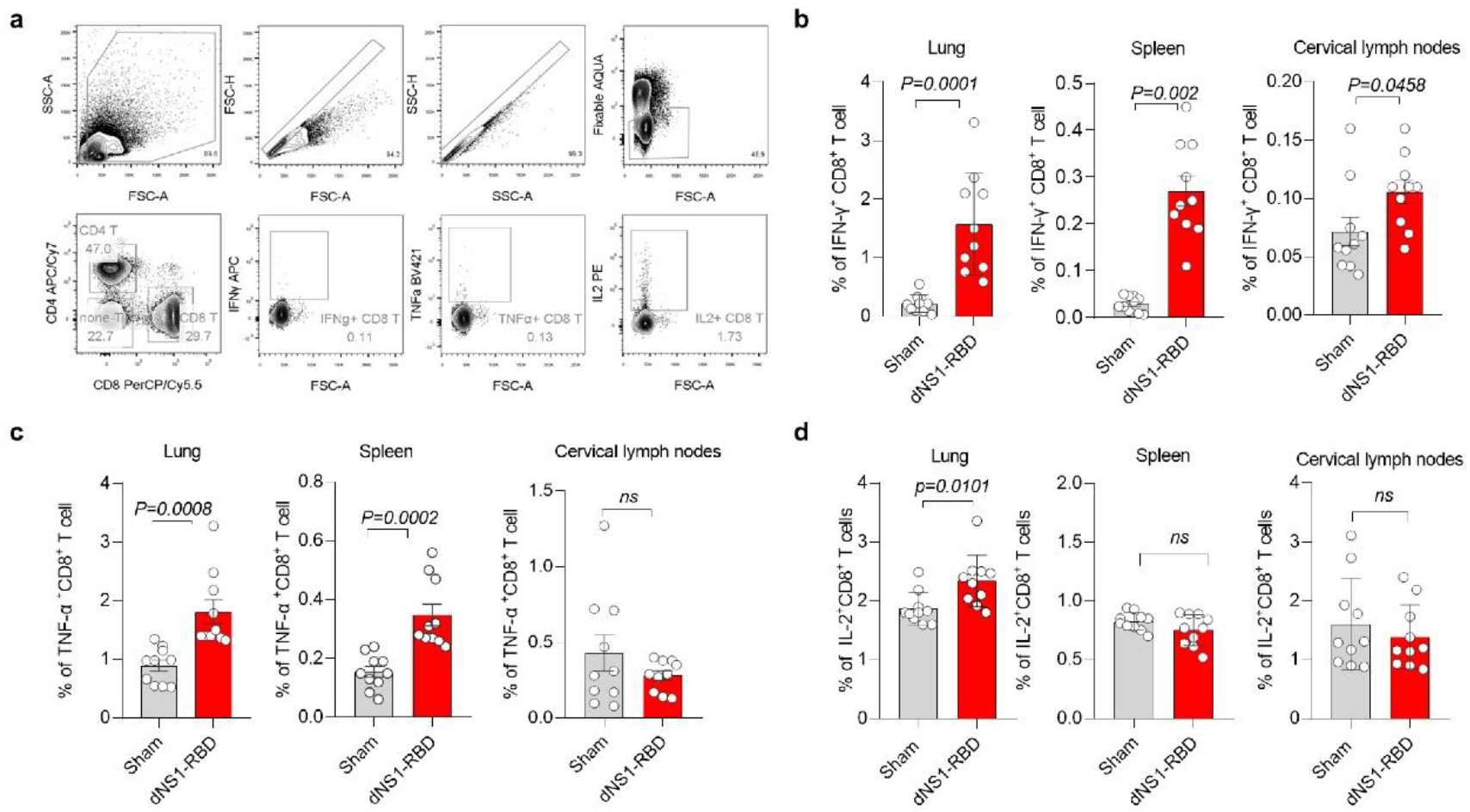
Gating strategy and profiling of CD8 T cells in dNS1-RBD-inoculated mice. **a** First, doublets were eliminated in the FSC-A *vs.* SSC-A plot, FSC-A *vs.* FSC-H plot and SSC-H *vs.* SSC-A plot. Then, live cells were selected by gating for fixable AQUA-positive and low FSC events. T cells were stratified into CD8 T cells. Boundaries defining positive and negative populations for intracellular markers were set based on nonstimulated control samples. **b-d** Intracellular cytokine staining assays for IFN-γ^+^CD8^+^ T cells (**b**), TNF-α^+^CD8^+^ T cells (**c**), and IL-2^+^CD8^+^ T cells (**d**) among the pulmonary lymphocytes, splenic lymphocytes or lymph node lymphocytes in response to the RBD peptide pool. Data are the mean ± SD; ns, not significant (P > 0.05); significance was determined by two-tailed Student’s t-test.

Serum samples and bronchoalveolar lavage fluid (BAL) were also collected 14 days after primary or booster immunization, and RBD-specific sIgA or IgG responses were evaluated by ELISA (Fig. 6g). The levels of RBD-specific sIgA and IgG titers increased significantly after boost immunization and peaked at 28 days post immunization, with all mice seroconverting. Whereas vaccines can induce the production of moderate levels of RBD-specific sIgA and IgG, the neutralizing activity of the induced antibodies was below the limit of detection (data not shown).

Overall, these data suggest that dNS1-RBD vaccination rapidly elicits vigorous and long-lived local innate and adaptive immune responses in the local respiratory tract that confer protection against SARS-CoV-2 infection (Fig. 6h).

## Discussion

To date, all COVID-19 vaccines approved are administered by intramuscular injection to elicit the production of primarily serum neutralizing antibodies and systemic T cell responses to fight against SARS-CoV-2 infection ^9^. However, intramuscular vaccines induce poor local immunity in the respiratory tract, which is the primary infection site for SARS-CoV-2^21^. It is evident that these vaccines are protective of severe diseases, however, breakthrough infections among vaccinated individuals are common ^11,12,33,34^. How to achieve more effective prevention of infection or transmission has become extremely important in the ongoing response to the COVID-19 pandemic.

One solution is to enhance the local immunity in the respiratory tract. Cold-adapted, live attenuated intranasal influenza vaccines have been used for more than a decade and shown to be effective to seasonal influenza, in particular among young children ^35^. Based on this concept, we have developed a live attenuated influenza vector (dNS1) by deleting viral immune modulator, the NS1 protein, from viral genome and identified adaptative mutations to support virus replication in eggs or MDCK cells which are commonly used for vaccine production. Using this dNS1 vector, we inserted the RBD gene of SARS-CoV-2 into the deleted NS1 site and made an influenza viral vector vaccine for COVID-19 (dNS1-RBD). This vaccine system has a few unique advantages which are immunogenic due to the lack of the NS1 which is a strong immune antagonist; it is extremely safe for use in all age groups; similar to the intranasal influenza vaccines, it is used intranasally to specifically induce mucosal immunity in the respiratory tract.

Our data showed that intranasal immunization of this dNS1-RBD vaccine is able to induce rapid protective and long-lasting immunities in hamsters when immunized hamsters were challenged 1 day or 7 days after single-dose vaccination or 9 months after booster vaccination. The protective immune response largely mitigated the lung pathology, with no apparent loss of body weight, caused by either the prototype-like strain or beta variant, suggesting cross-protective properties of this vaccine. To test if this vaccine can be used as rapid response to a sudden outbreak of SARS-CoV-2, we found it still renders protection when the hamster was vaccinated 24 h after challenge. Therefore, it is conducive to ring vaccination for the rapid establishment of protective immunity in high-risk populations in sporadic and epidemic infection areas. This study demonstrates that nasal vaccines may offer an attractive alternative in fighting against the COVID-19 pandemic.

What is special about this vaccine is that it is effective in preventing the pathological changes caused by COVID-19 without producing obviously detectable neutralizing antibodies, which is different from traditional vaccines mainly based on neutralizing antibodies. We believe that there are at least four aspects of the protective immune mechanism based on the current data. (i) Previous studies have reported that LAIVs induce the innate immune response in the nasal epithelium in animals, which not only is crucial for viral clearance and attenuation but also may play an important role in the induction of a protective immune response ^36^. In this study, we also observed the activation and secretion of antiviral cytokines and chemokines in lung tissue from vaccinated mice and correlated their production with rapid protection in hamsters. (ii) We believe that robust and local RBD-specific T cell responses should contribute to providing effective protection against SARS-CoV-2 infection ^37^. Considering resident memory CD8^+^ T cells, which are thought to provide long-lasting and broad-spectrum immune protection for LAIVs^38^, our data suggest that dNS1-RBD has the potential to confer long-lasting protective immunity, particularly around the bronchoalveolar space and lungs. Consistently, the hamster challenge results showed that dNS1-RBD conferred persistent protection against both the prototype-like strain and beta variant at 6 months after vaccination. (iii) Regarding the humoral immune response, RBD-specific IgA in BAL fluid and IgG in serum could be detected at a moderate level, which may effectively inhibit SARS-CoV-2 in the upper airways and nasal passages ^39^. (iv) Activated macrophages and myeloid DCs induced by dNS1-RBD, which are related to the short-term innate immune response, have recently been thought to exhibit memory characteristics through epigenetic reprogramming mechanisms, namely, trained immunity, which can exhibit faster and stronger immune responses to pathogen infections over a certain period of time ^31,40^. In addition, because the infected target cells of dNS1-RBD highly overlap with those of SARS-CoV-2, including both upper respiratory epithelial and alveolar epithelial cells^41–43^, the vaccine may also reshape the local anti-infection response of the respiratory tract through a mechanism similar to trained immunity. The contribution and mechanism of this effect in the protective immunity of the vaccine against respiratory tract infection deserve further study and exploration in the future.

LAIVs for intranasal administration were first licensed in the Soviet Union in 1987, in the US in 2003, and in Europe in 2012 and have a proven record of efficacy over decades of use ^35,44,45^. However, the immune response to LAIVs is multifaceted and does not necessarily involve a serum antibody response; LAIVs have been licensed on the basis of efficacy trials that measure protection rather than correlates of protection. A human challenge trial of LAIVs (FluMist^®^) also suggested that a low antibody response was not directly associated with low protective efficacy. In that study, the virus challenge results indicated that the LAIV had the equivalent and even improved efficacy of trivalent inactivated vaccine (TIV), while a higher seroresponse rate induced by the TIV ^20^. In general, the induction of the production of mucosal antibodies and a local T cell response by FluMist^®^ was similar to those induced by dNS1-RBD in adults (unpublished data).

Data from three earlier-phase clinical trials involving 1084 naïve adults showed that dNS1-RBD is very well tolerated and immunogenic in inducing production of mucosal IgA, systemic T cell responses and IgG against the RBD of SARS-CoV-2 (unpublished data). Undoubtedly, a phase III clinical trial conducted in COVID-19 epidemic countries is essential to finally determine the efficacies of dNS1-RBD against pivotal clinical outcomes associated with SARS-CoV-2 infection in humans, which is expected to be initiated soon (ChiCTR2100051391).

Thus, dNS1-RBD, an intranasally delivered vaccine candidate based on a live attenuated influenza virus vector, is unique for offering very rapid and prolonged broad protection against SARS-CoV-2 infection through inducing comprehensive local immune responses in the respiratory tract and might be a very promising vaccine which could fill the gap of current intramuscular vaccines. Further studies should be conducted to understand the unique immune activation and protection mechanism of intranasal immunization dNS1-RBD for SARS-CoV-2 in humans.

## MATERIALS and METHODS

### Cell cultures

All cell lines were obtained from ATCC. Human embryonic kidney cells (293T), African green monkey kidney epithelial cells (Vero E6), and Madin-Darby canine kidney cells (MDCK) were maintained in DMEM-high glucose (Sigma Aldrich, USA) supplemented with 10% low endotoxin FBS (Cegrogen Biotech, Germany) and penicillin-streptomycin.

### Construction of plasmids

The RBD segment of SARS-CoV-2 (GenBank accession number MN908947) was codon optimized for eukaryotic expression system and constructed by overlapping primers with the B2M signal peptide at the 5’ end and the foldon motif with the V5 tag at the 3’ end. The sequence encoding the RBD segment was then cloned into the NS1 deletion plasmid pHW2000-DelNS1 as described previously.

### Generation and passage of dNS1-RBD viruses

Eight pHW2000 plasmids containing the DelNS1 segment and the other seven influenza virus genomic segments, together with an NS1 expression plasmid, pCX-CA04-NS1-Flag, which derived from the parental influenza virus A/California/04/2009(H1N1) (GenBank: MN371610.1-371617.1), were transfected into 293T cells and incubated overnight at 37°C. The DNA mixture was removed, and Opti-MEM supplemented with 1 μg/mL TPCK-treated trypsin (Sigma) was added. Viral supernatant was collected 72 h later, designated dNS1-RBD passage 0 virus, and was subsequently passaged in MDCK cells at 33°C. The supernatant was harvested 48h post transfection when most of the cells showed signs of cytopathic effect (CPE). Infectious virus titers (PFU/mL) were determined by plaque assay on MDCK cells. The vaccine was further purified from supernatants by ultrafiltration, size-exclusion chromatography and then ion-exchange chromatography.

For the rescued viruses, deletion of the NS1 gene and insertion of the RBD gene were confirmed by reverse transcription-PCR (RT-PCR) using NS-specific and RBD-specific primers. Total RNA from virus supernatants was extracted using the PureLink™ Viral RNA/DNA Mini Kit (Thermo Fisher) according to the manufacturer’s protocol and then converted to cDNA by SuperScript® III Reverse Transcriptase (Invitrogen). The cDNA was then subjected to RT-PCR using primers and probes that are specific to the target sequence (RBD Forward-ACATTGGCCACCATGTTCACTGTAGAAAAAGGAAT; RBD Reverse- TTCCGGAATATAGCCGAAGTTGAAATTGACACATT; NS Forward- CCGAAGTTGGGGGGGAGCAAAAGCAGGGTGACAAAAACATA; NS Reverse- GGCCGCCGGGTTATTAGTAGAAACAAGGGTGTTTTTTATC, respectively). RT-PCR was performed under the following reaction conditions: 94°C for 2 min, followed by 35 cycles of 98°C for 15 s, 55°C for 30 s and 68°C for 90 s, and 68°C for 10 min. The presence of inserted sequences in generated vaccine virus were further confirmed by Sanger sequencing.

### Growth kinetics

MDCK cells seeded in 24-well plates were infected with viruses at the indicated multiplicity of infection (MOI). After 1 h of adsorption, the viral supernatant was removed, and the cells were washed twice with phosphate-buffered saline (PBS). DMEM containing 1 μg/ml TPCK-treated trypsin was added, and the cells were incubated at the indicated temperature. Supernatants were collected at different time points, and titers were determined by plaque assay.

### Plaque assay for dNS1-RBD viruses

Viruses were 10-fold serially diluted, added to confluent MDCK cells in 6-well plates and then incubated at 37°C for 1 h. The supernatant was removed, and the cells were washed twice with PBS and then overlaid with 1% MEM agarose containing 1 μg/ml TPCK-treated trypsin. The plates were incubated at 33°C for 72 h and then fixed with 4% PBS–buffered formaldehyde solution for at least 1 h. Plaques were visualized by staining with 1% crystal violet solution.

### Western blot

MDCK cells were cultured and infected with dNS1-RBD virus as described above. 36 hours later, cell lysates were harvested using modified NEP cell lysis buffer. Proteins were separated on a 10% gel, and then following transfer, blots were incubated with an anti-influenza A NP protein antibody 19C10 generated by our laboratory (1:1000) and anti-V5 tag antibody (Thermo,1:5000) and visualized with horseradish peroxidase (HRP)-conjugated anti-mouse IgG (Invitrogen, 1:5000).

### Immunofluorescence imaging

For direct visualization of the expression of HA and RBD, MDCK cells were seeded at 2×10^4^ cells per well in CellCarrier-96 Black plates and then infected with dNS1-RBD, CA04-dNS1 and CA04 WT at an MOI of 1. PBS was used as a negative control. After 72 h, the cells were fixed with 2% paraformaldehyde in PBS for 15 min in the dark. The cells were then permeated by the addition of 0.3% Triton X-100 in PBS (PBST) for 10 min at room temperature and blocked with 2% BSA. The plates were incubated with a DyLight 488-labeled mAb against 6G9-488 (anti-HA; 1:100 dilution) and DyLight 650-labeled mAb against R4D11 (anti-RBD; 1:100 dilution) generated by our laboratory at 37°C for 60 min, and the assay plates were washed three times with PBS. Cell nuclei were labeled with DAPI. The images were acquired on an Opera Phenix using a 63× water immersion objective.

### Vaccine formulation

The vaccine dNS1-RBD was prepared on a large scale at Beijing Wantai Biological Pharmacy Enterprise Co., Ltd., Beijing, China. After rounds of passage and amplification with the cell factory based on the MDCK cell line, purified dNS1-RBD virions were mixed with virus protectant, which contained carbohydrates, amino acids and human albumin, etc., and were preserved at −15°C. Based on the ELISA results using a sandwich assay with anti-RBD monoclonal antibodies on both sides (Wantai, Beijing, China) and plaque assay results, serial passages 1 to 10 of purified vaccines were confirmed to be stable under current vaccine manufacturing conditions.

### Vaccine safety evaluation

The safety of the potential SARS-CoV-2 vaccine, dNS1-RBD was evaluated in BALB/c mice and ferret. BALB/c mice were intranasally inoculated with 10^5^-10^7^ PFU of dNS1-RBD and CA04-WT under isoflurane anesthesia and monitored daily for morbidity and mortality for 14 days post inoculation. Vaccines were diluted in 1640 media to a final 50 μL volume and administered bilaterally for BALB/c mice. Animals that lost more than 25% of their initial body weight were euthanized in accordance with our animal ethics protocol. Ferret studies were performed at JOINN Labs (Suzhou). Two groups of ferrets (5 female and 5 male ferrets in the vaccine group and 3 female and 3 male ferrets in the control group) were immunized intranasally with a single-dose 1×10^6^ PFU of dNS1-RBD and CA04-WT virus respectively diluted in 1640 media to a final 500 μL volume. Datasets of the safety-related parameters were collected during and after immunization, including clinical observations, body weight, and body temperatures. Viral loads were detected for throat swabs and nasal washes collected at days −1, 1, 3, 5, and 7 post-inoculation by RT-PCR. Histopathological evaluations in lungs from two groups of ferrets were conducted at day 8. Lung tissues were collected and stained with hematoxylin and eosin. Six to eight-weeks-old, female BALB/c mice and five to six-months-old, male and female ferret were used throughout this study.

### Immunization and infection of mice

BALB/c mice were immunized intranasally with 50 μL containing 1×10^6^ PFU of the vaccine prepared as indicated above under isoflurane anesthesia, while the control group was administered CA04-WT or CA04-dNS1 virus. For antibody response evaluation, all groups of BALB/c mice (6 animals in each group) were vaccinated by a prime-boost regimen (days 0 and 14), and blood was collected via retro-orbital bleeding before each immunization and 14 days after the second injection, followed by a binding assay to analyze vaccine immunogenicity.

For innate immune response analyses, C57BL/6 mice (5 animals in each group) were vaccinated with a single dose and sacrificed 1 day after vaccination. For cellular immune response analyses of PBMCs, splenic lymphocytes, pulmonary lymphocytes and lymph node cells, C57BL/6 mice (6-8 weeks old) were immunized intranasally with 1×10^6^ PFU of the vaccine by the one-dose or two-dose regimen as described above (10 animals in each group). Then, splenic lymphocytes, pulmonary lymphocytes and lymph node cells (6 animals in each group) were collected on day 28 of a prime-boost regimen with a 2-week interval for intracellular cytokine staining (ICS) measurements.

### Immunization and infection of hACE2-KI/NIFDC mice

hACE2-KI/NIFDC mice (8-10 weeks old) were divided into three groups and treated intranasally with 1×10^6^ PFU of the vaccine by gently adding 50 μL droplets of virus stock for the vaccine-immunized group (5 animals) at two time points (days 0 and 14), and then, the vaccine-immunized group and unvaccinated group (3 animals each) were challenged with 1×10^4^ PFU of SARS-CoV-2 by the intranasal route 30 days post immunization. Datasets of safety-related parameters were collected throughout the whole assay, including clinical observations and body weight. Three lung lobes of all nine euthanized hACE2-humanized mice were collected at 4 days post challenge and used for RT-PCR, plaque and histopathological assays.

### Immunization and infection of hamsters

Hamsters (male:female=1:1) were vaccinated with the indicated amount of the vaccine. All hamsters received 100 μL of vaccine per dose via the intranasal route. At the indicated time after vaccination, the hamsters were further evaluated by direct contact challenge of SARS-CoV-2. Two strains were used in this study: the prototype-like virus AP8 (hCoV-19/China/AP8/2020; GISAID accession number: EPI_ISL_1655937) and the beta variant AP100 (hCoV-19/China/AP100/2021; GISAID accession number: EPI_ISL_2779638). Virus-carrying hamsters (donors) were preinfected via inoculation of 1×10^3^ PFU of SARS-CoV-2 through the intranasal route. Each donor was transferred to a new cage and cohoused with four vaccinated or unvaccinated control animals. One day after cohousing, donors were isolated from the cage, and the other hamsters were further observed. The hamsters were fed a daily food amount of 7 g per 100 g of body weight. The weight changes and typical symptoms (piloerection, hunched back, and abdominal respiration) in hamsters were recorded daily after virus inoculation or contact. Hamsters were sacrificed for tissue pathological and virological analyses on day 5 after virus challenge. The virus challenge studies were performed in an animal biosafety level 3 (ABSL-3) facility.

### Anti-RBD IgA measurements

Bronchoalveolar lavage (BAL) was collected on control-infected and vaccine-infected mice. Mice were euthanized, and a short needle insulin syringe (BD, USA) was inserted gently into the lumen of the exposed trachea. The lungs were then lavaged with two separate 1-mL washes of sterile normal saline. The RBD-specific IgA titer of BAL samples was next evaluated by ELISA as described above with Goat anti-mouse IgA alpha chain-HRP (Abcam, 1:3000).

### Anti-RBD IgG measurements

RBD-specific antibody titers in serum samples collected from immunized animals with 1×10^6^ PFU of vaccine were determined by indirect ELISA. Ninety-six-well microtiter plates were coated with 200 ng of purified RBD protein which was generated and expressed in 293F from the codon optimized RBD sequence of SARS-CoV-2 spike protein (GenBank accession number MN908947) individually at 4°C overnight and blocked with 2% BSA for 2 h at 37°C. Diluted sera (1:100) were successively diluted in a 2-fold series and applied to each well for 1 h at 37°C, followed by incubation with goat anti-mouse, anti-hamster or anti-human antibodies conjugated with HRP for 1 h at 37°C after 3 washes. The plate was developed using TMB, followed by the addition of 2M H_2_SO_4_ to stop the reaction, and read at 450/630 nm by ELISA plate reader for final data acquisition.

### ELISPOT assay

ELISPOT assays were performed using mouse IFN-γ ELISpot plates (DAKEWE). Ninety-six-well ELISpot plates precoated with capture antibody were blocked with RPMI-1640 for 10 min at room temperature. Briefly, a total of 10^6^ cells per well from C57BL/6 mouse spleen, lymph nodes, lung or PBMCs immunized with 1×10^6^ PFU of vaccine were plated into each well and stimulated for 20 h with pooled peptides of RBD of wild type SARS-CoV-2 or variants (15-mer peptide with 11 amino acids overlap, cover the spike, Genscript). The spots were developed based on the manufacturer’s instructions. PBS and cell stimulation cocktails from the kit were used as negative and positive controls, respectively. Spots were scanned and quantified by an ImmunoSpot CTL reader. Spot-forming units (SFUs) per million cells were calculated by subtracting the negative control wells.

### Intracellular cytokine staining assay

The expression of phenotypic markers, activation markers, and cytokines was evaluated using flow cytometry for T cells, B cells, and monocytes/macrophages in single-cell suspensions from tissues. The cells were stained with murine antibodies for phenotype and activation (CD4 [clone GK1.5, APC/Cy7], CD8 [clone 53-6.7, PerCP/Cy5.5], CD11b [clone M1/70, PE], CD11c [clone N418, BV421], CD49b [clone dx5, FITC], MHC2 [clone M5/114.15.2, APC], Ly-6C [clone HK1.4, APC-Cy7], Ly-6G [clone 1A8, BV605], CD62L [clone MEL-14, APC-Cy7], CD103 [clone 2E7, PE], CD69 [clone H1.2F3, BV421], CD44 [clone IM7, APC], CD80 [clone 16-10A1, FITC], CD86 [clone GL-1, PE-Cy7]) and cytokine expression (IL4 [clone 11B11, BV421], IL2 [clone JES6-5H4, PE] and IFN-γ [clone XMG1.2, APC]), and a LIVE/DEAD® Fixable Aqua Dead Cell Stain Kit was also used. For RBD-specific T cell assays, each sample was stimulated with pooled spike peptides (1 μg/well) in a U-bottom plate and incubated at 37°C for 18 h. After incubation, 0.12 μL of protein transport inhibitors (BD GolgiPlug^TM^, BD Biosciences) in 20 μL of 10% FBS/RPMI 1640 medium was added to each well, and the plate was incubated at 37°C for 6 h. Then, the cells were washed once with 2% FBS/PBS and further stained with labeled antibodies. After incubation at 4°C for 30 min, the cells were washed once with 2% FBS/RPMI 1640 medium and passed through a 0.22 μm filter. The cells were transferred to 5-mL round-bottom tubes and analyzed by a BD LSRFORTESSA X-20 system. The data were analyzed by FlowJo V10.6.0 and GraphPad Prism 9.

### Measurements of cytokine and chemokine levels

Lung homogenate samples were prepared for analysis with ProcartaPlex Multiplex Immunoassay, a mouse cytokine/chemokine magnetic bead panel (36-plex, Thermo Fisher, MA, USA), following kit-specific protocols. Analytes were quantified using a Magpix analytical test instrument using a standard curve derived from recombinant cytokine and chemokine standards, which utilizes xMAP technology (Luminex Corp., Austin, TX) and xPONENT 4.2 software (Luminex). The results were expressed as ng/mL.

### SARS-CoV-2 and dNS1-RBD RNA quantification

Viral RNA levels in the lungs of challenged hamsters were detected by quantitative RT-PCR. Briefly, for quantification of viral levels and gene expression after challenge or passage experiments, RNA was extracted from homogenized organs or cultured cells using a QIAamp Viral RNA Mini Kit (Qiagen) according to the manufacturer’s instructions. Hamster tissue samples were homogenized by TissueLyser II (Qiagen, Hilden, Germany) in 1 mL of PBS. Subsequently, viral RNA quantification was conducted using a SARS-CoV-2 RT-PCR Kit (Wantai, Beijing, China) by measuring the copy numbers of the N gene, while CA4-dNS1-nCoV-RBD was quantified with primers targeting the RBD and NS genes.

### SARS-CoV-2 titration assay

Live virus titers in homogenized lung tissues and cell cultures were measured by the standard TCID_50_ method in Vero E6 cells seeded in 96-well plates. In brief, the samples were serially diluted, added to the 96-well plates and incubated with the Vero E6 cells for one hour. Three days after incubation, the cytopathic effects were observed and used to calculate the viral titers.

### Histopathology

The lung tissues from challenged hamsters were fixed with 10% formalin for 48 h, embedded in paraffin and sectioned. Next, the fixed lung sections were subjected to hematoxylin and eosin (H&E) staining. Immunohistochemical staining was performed by using a mouse monoclonal anti-SARS-CoV-2 N protein antibody. Whole-slide images of the lung sections were captured by an EVOS M7000 Images System (Thermo Fisher).

### Statistical analysis

Statistical significance was assigned when P values were < 0.05 using GraphPad Prism 8.0 (GraphPad Software, Inc.). Viral titers and RBD-specific IgG titers were analyzed after log-transformation. The bars in this study represent the mean ± SD or median (interquartile range, IQR) according to data distribution. The number of animals and independent experiments that were performed are indicated in the figure legends. Student’s t-test (two groups) or one-way analysis of variance (ANOVA) (three or more groups) was used for comparison of normally distributed continuous variables. For nonnormally distributed continuous variable comparisons, the Mann-Whitney U test (two groups) or Kruskal-Wallis test (three or more groups) was used. Two-way repeated-measures ANOVA was adopted for repeated data comparison. For multiple comparisons of three or more groups, Dunnett’s multiple comparison test was used.

### Ethics statements

All animals involved in this study were housed and cared for in an Association for the Assessment and Accreditation of Laboratory Animal Care (AAALAC)-accredited facility. All experimental procedures with mice, ferrets and hamsters were conducted according to Chinese animal use guidelines and were approved by the Institutional Animal Care and Use Committee (IACUC) of Xiamen University. The hACE2 and hamster studies were performed in an animal biosafety level 3 (ABSL-3) laboratory affiliated to the State Key Laboratory of Emerging Infectious Diseases, The University of Hong Kong.

## ACKNOWLEDGEMENTS

This work was supported by the National Program on Key Research Project of China (2020YFC0842600), the National Natural Science Foundation of China (82041038, 81871651, 81991491), and Science and Technology Major Program of Fujian Province (2020YZ014001).

## AUTHOR CONTRIBUTIONS

J.C., P.W., L.Y., Liang Z., Limin Z., H.Z., C.C., Y.C, Jinle H., J.J., Zen L., Junping H., L.C., Zicen L., Q.W., R.C., M.C., R.Q., X.W., J.M., M.Z., H.Y., Q.H., and G.W. performed experiments and analyzed data. X.Y., C.L., T.Z., J.Z., H.Z., Y.C., H.C. and N.X. designed and supervised the study. J.C., T.Z., J.Z., H.Z., Y.C., H.C. and N.X. wrote the manuscript. C.F., C.Z., X.L., Y.S., Q.Y., T.C., and T.W. participated in design and result analysis.

