## Supplementary material for "A live attenuated influenza virus-vectored intranasal COVID-19 vaccine provides rapid, prolonged, and broad protection against SARS-CoV-2 infection": Supplenmentary information

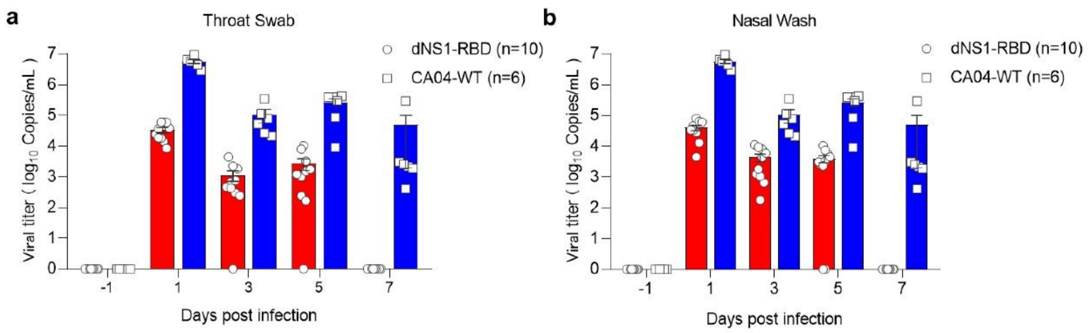


**Supplementary Fig. S1 Viral shedding of dNS1-RBD in ferrets. a-b**, Two groups of ferrets were immunized with a single dose of the dNS1-RBD (red) and CA04-WT (blue) vaccines through the intranasal route. Throat swabs (**a**) and nasal washes (**b**) of ferrets collected at days -1, 1, 3, 5, and 7 were assayed.

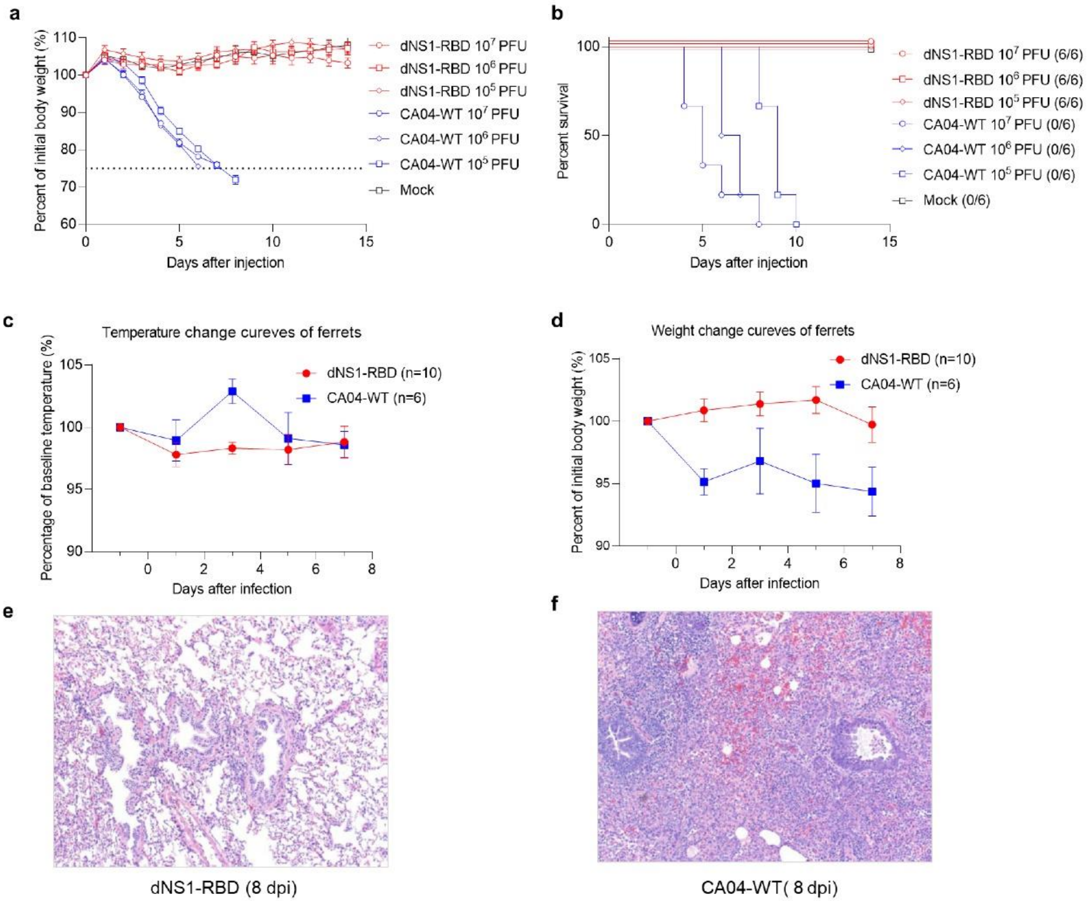


**Supplementary Fig. S2 Pathogenicity evaluation of dNS1-RBD in mice and ferrets.**

**a-b**, Body weight changes (**a**) and survival (**b**) of female BALB/c mice following intranasal administration of different doses of dNS1-RBD ($10^5$, $10^6$ and $10^7$ PFU) in comparison to those following intranasal administration of different doses of CA04-WT ($10^5$, $10^6$ and $10^7$ PFU). **c-d**, Two groups of ferrets intranasally administered a single-dose dNS1-RBD vaccine (n=10) or CA04-WT (n=6) showed attenuation of dNS1-RBD infection. (**c**) Body weight and (**d**) body temperature change of ferrets. **e-f**, Histopathological evaluations of the lungs from the two groups of ferrets at day 8 post administration. Lung tissues were collected and stained with hematoxylin and eosin. Inoculation with $10^7$ PFU of CA04-WT resulted in obvious influenza-like symptoms with fever, weight loss and pathological injury in lung tissues from ferrets.

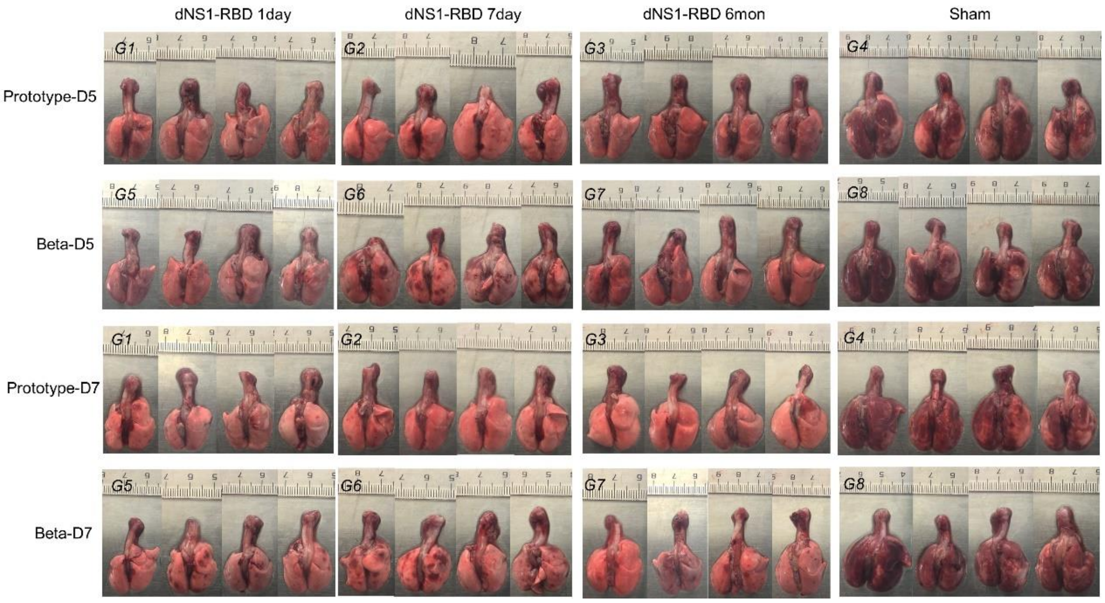


**Supplementary Fig. S3 Gross observations of lung tissues from hamsters in groups 1 to 8 as indicated in Figure 2 of the main manuscript.** The immunized hamsters were euthanized, and the lungs were isolated at 5 dpi and 7 dpi. Severe lung lesions, including consolidation and multifocal and diffuse hyperemia, were observed in G4 and G8 hamsters, while focal histopathological changes in a few lobes of the lung were observed in the vaccinated hamsters.

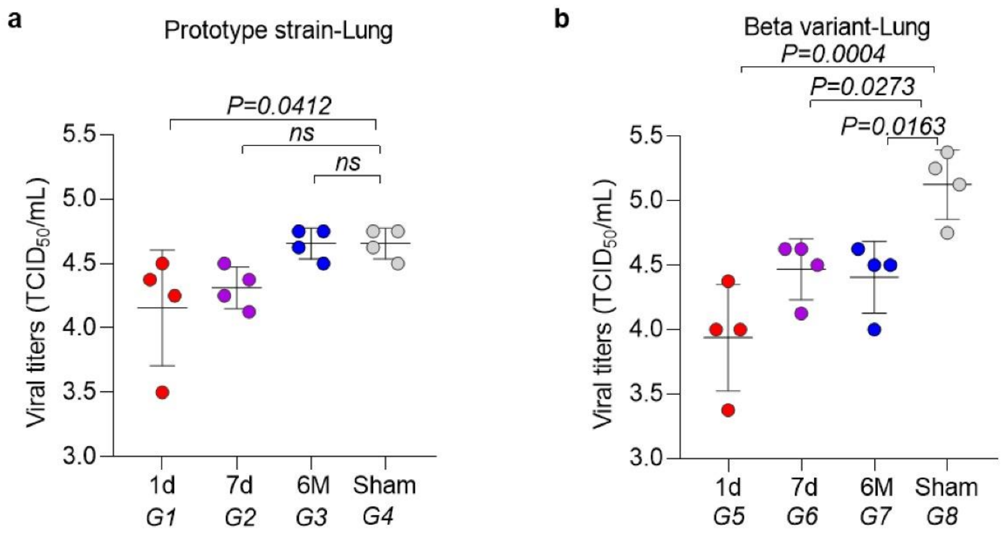


**Supplementary Fig. S4 Viral loads of lung tissue from challenged hamsters. a-b**, There were eight total experimental groups each containing eight hamsters (males:females=1:1). **a**, Groups 1, 2, 3 and 4 were challenged with the prototype SARS-CoV-2 strain; **b**, groups 5, 6, 7 and 8 were challenged with the beta variant. Groups 1 and 5 received a single dose of dNS1-RBD 1 day before challenge; groups 2 and 6 received a single dose of dNS1-RBD 7 days before challenge; groups 3 and 7 received two doses of dNS1-RBD at a 14-day interval 6 months before challenge; and groups 4 and 8 served as sham controls and were not treated. Viral loads of lung tissue obtained at 5 dpi from hamsters challenged by the prototype (**a**) or beta strain (**b**) were determined by TCID50 assay. Data are the mean ± SD; ns, not significant ($P > 0.05$); significance was determined by ordinary one-way ANOVA multiple comparison.

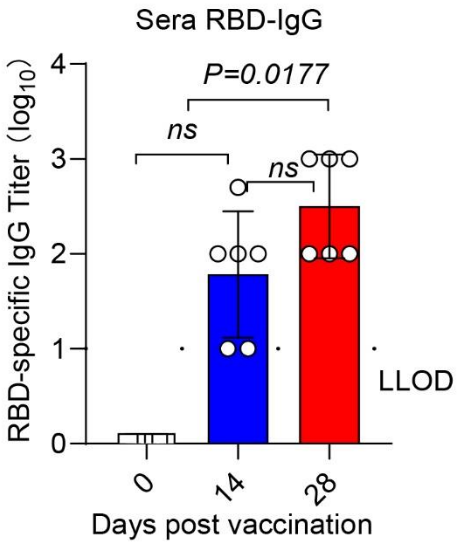


**Supplementary Fig. S5 Quantification of humoral response levels in hamsters.** RBD-specific IgG levels in serum were measured by ELISA for hamsters vaccinated twice, at day 0 and day 14. Data for the antibody analysis are presented as the geometric mean with the geometric SD from four independent experiments. LLOD-lower limit of detection. Data are the mean ± SD; ns, not significant ($P > 0.05$); significance was determined by ordinary one-way ANOVA multiple comparison.

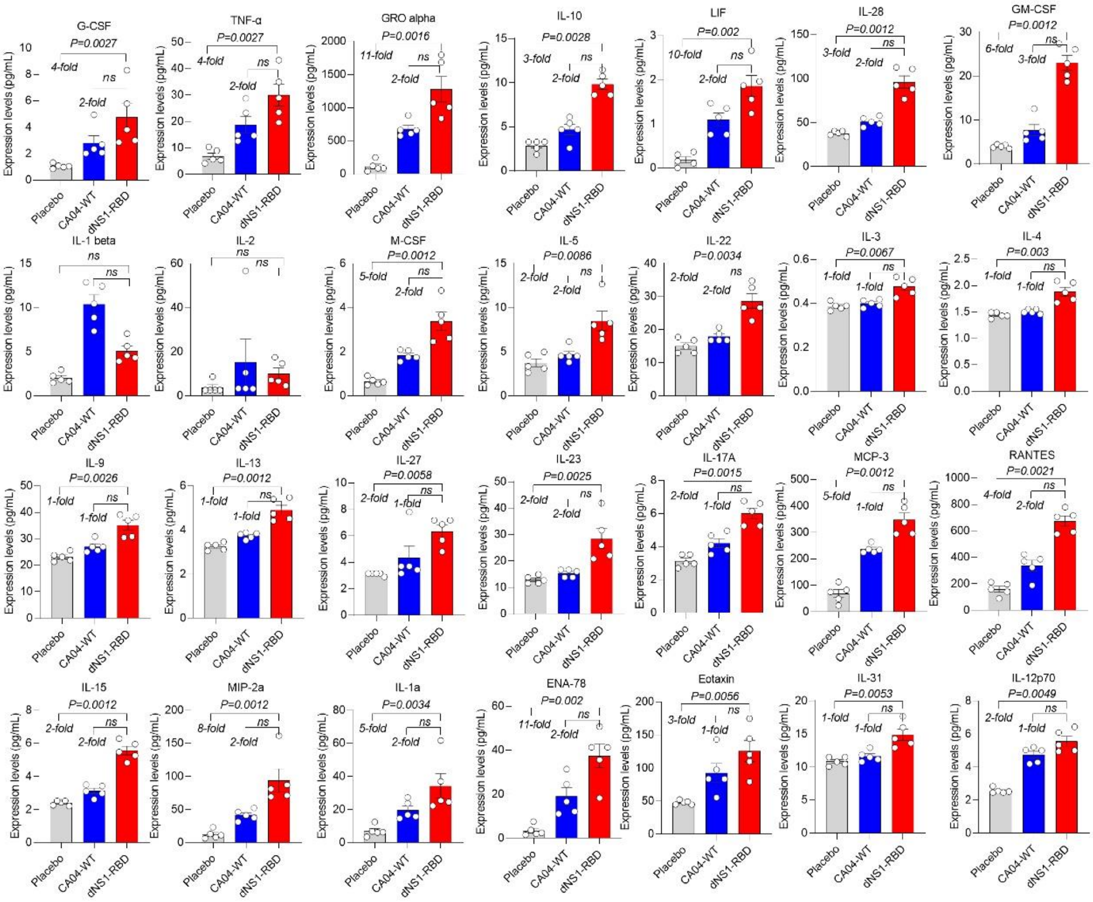


**Supplementary Fig. S6 Quantification of cytokine and chemokine expression levels.** Lung homogenates from BALB/c mice vaccinated with dNS1-RBD one day prior to sacrifice were collected for quantification of the other 28 cytokine and chemokine expression levels except for IL-6, IL-18, IFN-γ, IFN-α, MCP-1, IP-10, MIP-1α, and MIP-1β by ProcartaPlex immunoassays. The data are expressed as ng/mL. Data are the mean ± SD; ns, not significant ($P > 0.05$); significance was determined by ordinary one-way ANOVA multiple comparison.

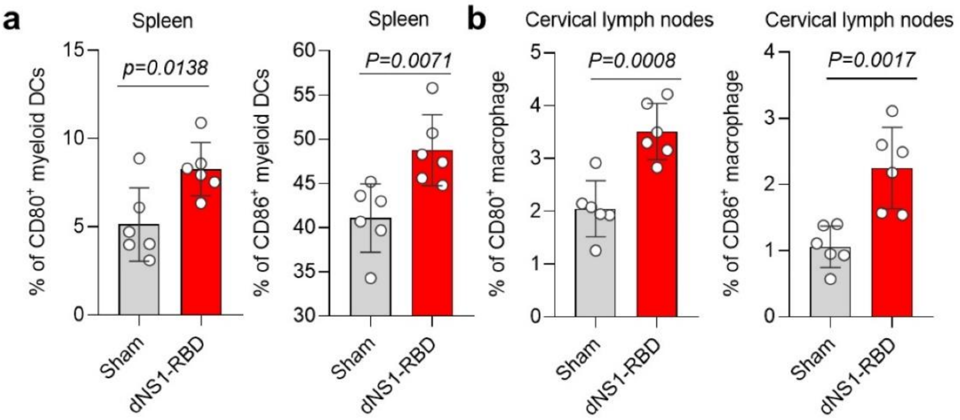


**Supplementary Fig. S7 Activation and differentiation of various innate immune cells. a-b**, CD80 and CD86 expression on antigen-presenting cells from immunized C57BL/6 mouse spleens (**a**) and cervical lymph nodes (**b**) at 14 days after vaccination. Data are the mean ± SD; significance was determined by two-tailed Student's t-test.

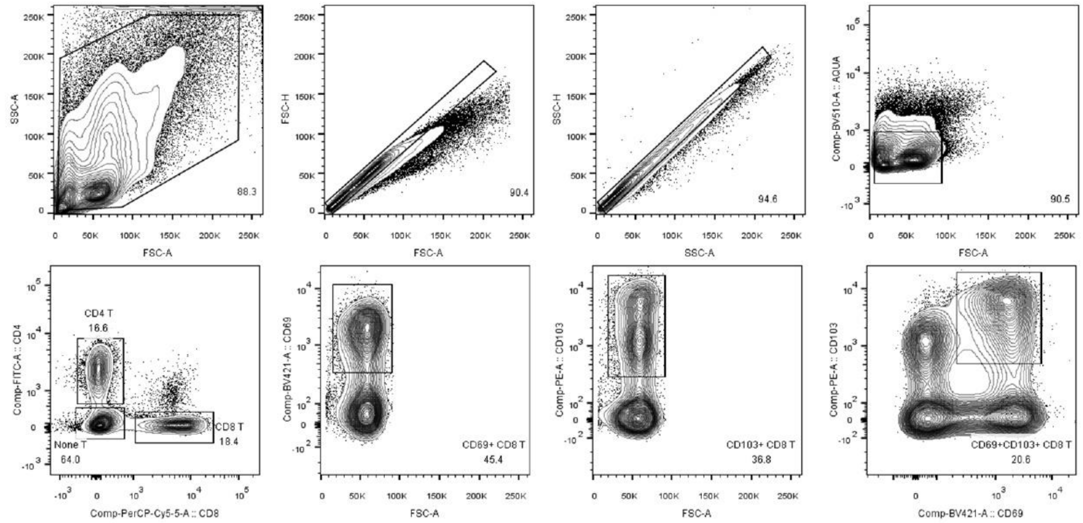


**Supplementary Fig. S8 Gating strategy for identifying pulmonary T cells expressing CD69⁺ and CD103⁺ in dNS1-RBD-infected mice.** Mice inoculated intranasally with dNS1-RBD. Representative flow cytometry profiles are from mice harvested at different times.
